# Striatal habit system drives behavioral rigidity in PTSD

**DOI:** 10.64898/2026.09.17.752357

**Authors:** Krystian B. Loetscher, D.T. Nguyen, Sanghoon Kang, John H. Krystal, Stephanie M. Groman, Elizabeth V. Goldfarb

## Abstract

Core symptoms of posttraumatic stress disorder (PTSD) like safety seeking and avoidance have long been described as habitual. Although widespread theoretical accounts posit a relationship between these behaviors and striatal-mediated habits, or rigid stimulus-response (S-R) associations, this idea has yet to be tested. Here, we implement a new task design to investigate how S-R associations are acquired, how flexibly they can be remapped, and how sensitive they are to changes in outcome value for individuals with PTSD as compared to individuals who experienced trauma without developing PTSD (trauma-exposed controls; TC). We find that individuals with PTSD, compared to TC, exhibit several hallmarks of habitual behavior, including more rigid habit-like perseveration following response remapping, and, with more severe symptoms, continued perseveration following outcome devaluation. We also observe that the PTSD group acquired these S-R associations more slowly. Using functional magnetic resonance imaging, we find significant putamen engagement when learning S-R associations that was especially pronounced in PTSD and predicted later perseveration. Finally, computational models indicate that the PTSD group leveraged gradual trial-by-trial updating to learn and later update associations, whereas the TC group leaned more on a stable, consistent choice strategy, both of which explained later behavioral rigidity. In addition to distinguishing individuals with PTSD, these habit-related processes were also more pronounced for individuals with more severe symptoms. Together, these results provide the first empirical evidence for the role of habits in PTSD.

## Introduction

The observation that rigid habits can develop after trauma has a long history in the clinic, rodent models, and even in literature. Consider the indelible image of Lady Macbeth repeating the motion of scrubbing her clean hands after the murder of the king. Clinically, post-combat stress has been posited to provide a “rich culture medium for the operant conditioning of what may be lasting symptoms”^1^. Nevertheless, empirical evidence for the relationship between habits and posttraumatic stress disorder (PTSD) is lacking. Although contemporary models of PTSD emphasize neural mechanisms of encoding and retrieving fear^2,3^, many persistent symptoms cannot be explained by these frameworks^4^. Indeed, avoidance behaviors across species have long proven resistant to interventions involving fear reduction^5^. Understanding the neural mechanisms underlying such symptoms (like rigid avoidance of trauma cues and safety seeking behavior) is essential to improving clinical interventions^6–9^. Here we present the first empirical test of the relationship between habits and PTSD.

Habits are characterized by rigid stimulus-response (S-R) associations that are gradually acquired and difficult to change. These responses are both insensitive to changes in outcome value (e.g. devaluation), and to response-outcome contingency changes (e.g., new response required for different outcomes) often manifesting as perseveration. At a neural level, habits are associated with the dorsolateral striatum (DLS; human putamen)^10–14^. These classic habit-related processes bear a striking resemblance to symptoms of PTSD that are rigid and resistant to change, namely avoidance behaviors (e.g. avoiding restaurants) and other arousal-reducing actions (e.g. scanning for exits). Like habits, these symptoms and related behaviors like cue-induced intrusions and perseverative thinking are also automatic and implicit (e.g. difficult to integrate into declarative memory)^15–17^. Nevertheless, the relationship between PTSD and habit lacks empirical support^15,18–20^.

There is a large body of evidence suggesting that PTSD may be associated with stronger expression of habitual behaviors. Consistent with the pivotal role of past stress in the etiology of PTSD, both recent and past stress lead to more reliance on S-R associations across species^21^, manifesting as decreased sensitivity to changes in outcome value and reduced flexibility in adapting behavior when circumstances change^22–25^. Even early life stress exposure has been associated with persistent habit-like behaviors in adulthood^26–28^. At the neural level, extreme stress leads to long-lasting reorganization of fronto-striatal circuits, enhances amygdala-striatal connectivity, and increases dendritic arbors and complexity in the DLS^20,22,29–32^.

However, not all individuals who experience stress – even extreme stress – go on to develop PTSD. Recent theoretical and empirical work suggest that individuals with PTSD exhibit sustained elevation of arousal, orchestrated by the amygdala and, intriguingly, perhaps the DLS^19,33,34^. There is recent evidence that individuals with stronger post-trauma psychopathology show greater DLS responses to reward-predictive cues^35^. In individuals who develop PTSD, chronic hyperarousal may constitute a persistent internal state driving habit circuitry. For instance, heightened arousal states have been shown to favor habitual control in rodents; anxiogenic drugs, predator odor, and fear-conditioned stimuli all induce a shift towards DLS-mediated S-R behavior^34,36,37^. Emotional arousal observed in PTSD may also lead to more persistent habit-like behaviors over one month later^38^. Accordingly, several theories posit that hypervigilance – a state of persistently elevated arousal – may contribute to rigid habit-like behaviors^1,33,39^. Indeed, the DLS has also been implicated in hypervigilance in schizophrenia^40^ and vigilant attention more broadly^41^. Together, these results suggest that both long-lasting changes induced by stress and an enduring state of hypervigilance may drive maladaptive and persistent reliance on DLS-mediated habits.

Here we investigated the roles of behavioral, neural, and computational mechanisms associated with habit in PTSD. We hypothesized that individuals with PTSD would evince stronger habit expression as well as greater reliance on DLS (putamen) when forming these associations. Given previous work demonstrating altered prediction-error based learning following trauma, we also hypothesized that they would use different learning computations to build these rigid associations^42–44^.

To test these hypotheses, we developed a new computerized task in which participants learn to place images of different objects in one of two rooms, with correct responses yielding a stimulus-unique reward (a fruit image that adds to their monetary bonus; **Fig. 1a**). Placing an object into a room required a brief motor sequence to facilitate striatal chunking ^45,46^. Participants completed this task while undergoing functional magnetic resonance imaging (fMRI). Participants acquired S (object) – R (room selection) associations with probabilistic (80%) reinforcement ^45,47^, learning six interleaved S-R associations to reduce reliance on working memory ^48^ (**Fig. 1b**). This Learning phase (300 trials over 10 blocks) allowed us to quantify the formation of S-R associations and associated putamen engagement^12^. Next, with no information signaled to the participants, they were required to update their responses (i.e., room selection) for two of the six stimuli ^49,50^. This Reversal phase (300 trials over 10 blocks) allowed us to assess flexibility via response remapping. Finally, participants were instructed that two of six fruit outcomes were now ‘rotten’ and would detract from their monetary bonus, requiring participants to avoid earning those fruits. They then continued placing objects in different rooms without feedback. This Devaluation phase (30 trials over 1 block) provides a classic test of flexibility via sensitivity to changing outcome value^51^ (**Fig. 1c**).

**Figure 1.**
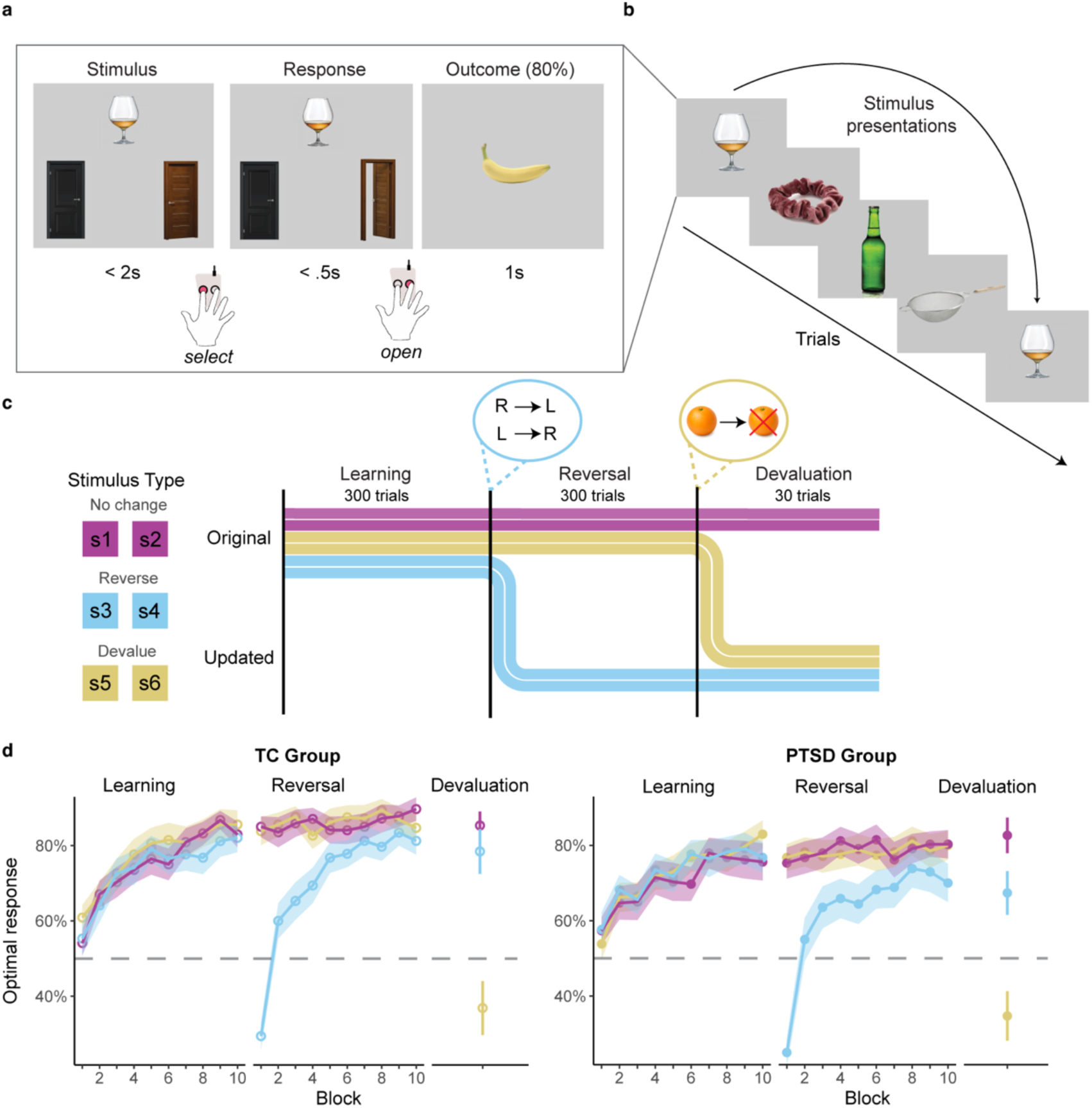
Task design and validation. **a.** Single trial. **b.** Trial sequence with randomly interleaved stimuli. **c.** Task phases. Participants learn 6 unique stimuli: 2 reverse in the Reversal phase, 2 have associated outcomes devalued in the Devaluation phase, and the remaining 2 remain unchanged. **d.** Mean optimal responding per stimulus type and phase in each group. Ribbons/error bars = ± 1 SEM.

## Results

### Both PTSD and TC groups successfully learn and update probabilistic associations

We first confirmed that participants in both groups could successfully learn and update probabilistic S-R associations (**Fig. 1d**, for demographics see Table S1). By the end of Learning, both groups made the optimal choice (e.g., 80% reinforced) more often than chance (TC: all *ts*(33) > 7.83, all *ps* < 0.001; PTSD: all *ts*(33) > 5.97, all *ps* < 0.001). By the end of Reversal, both groups updated their responses to stimuli with remapped responses (updated response vs chance; TC: *t*(31) = 9.99, p < 0.001; PTSD: *t*(33) = 4.49, *p* < 0.001) and maintained above-chance performance to stimuli with unchanged responses (unchanged and to-be-devalued stimuli; TC: all *ts*(31) > 12.22, *p* < 0.001; PTSD: all *ts*(33) > 7.87, all *ps* < 0.001). Following outcome devaluation, both groups showed successful value updating by selecting the devalued fruits significantly less often than valued fruits in the consumption test (valued vs devalued; TC: *t*(31) = 6.59, *p* < 0.001; PTSD: *t*(33) = 6.53, *p* < 0.001; Supplementary Fig. 1a). Furthermore, participants in both groups were significantly less likely to persist in their responses to stimuli paired with these devalued outcomes relative to unchanged stimuli with valued outcomes (TC: *t*(31) = 5.59, *p* < 0.001; PTSD: *t*(33) = 5.81, *p* < 0.001). Verifying that both groups attended to the task, we found comparable (low) numbers of skipped trials across phases between groups (*F*(1,66) = 1.86, *p* = 0.178) and above-chance memory for which outcomes were paired with each stimulus in both groups (all ps < 0.001; Supplementary Fig. 1b).

### Differences in learning between PTSD and TC groups

By the end of Learning, both PTSD and TC groups achieved similar performance (TC vs PTSD; *t*(66) = −0.88, *p* = 0.380; **Fig. 2a**). Examining each of the six S-R associations separately revealed no group differences in the number of associations learned (i.e., stimuli with above-chance performance; main effect group: *F*(1,66) = 0.014, *p* = 0.907, Supplementary Fig. 2a).

**Figure 2.**
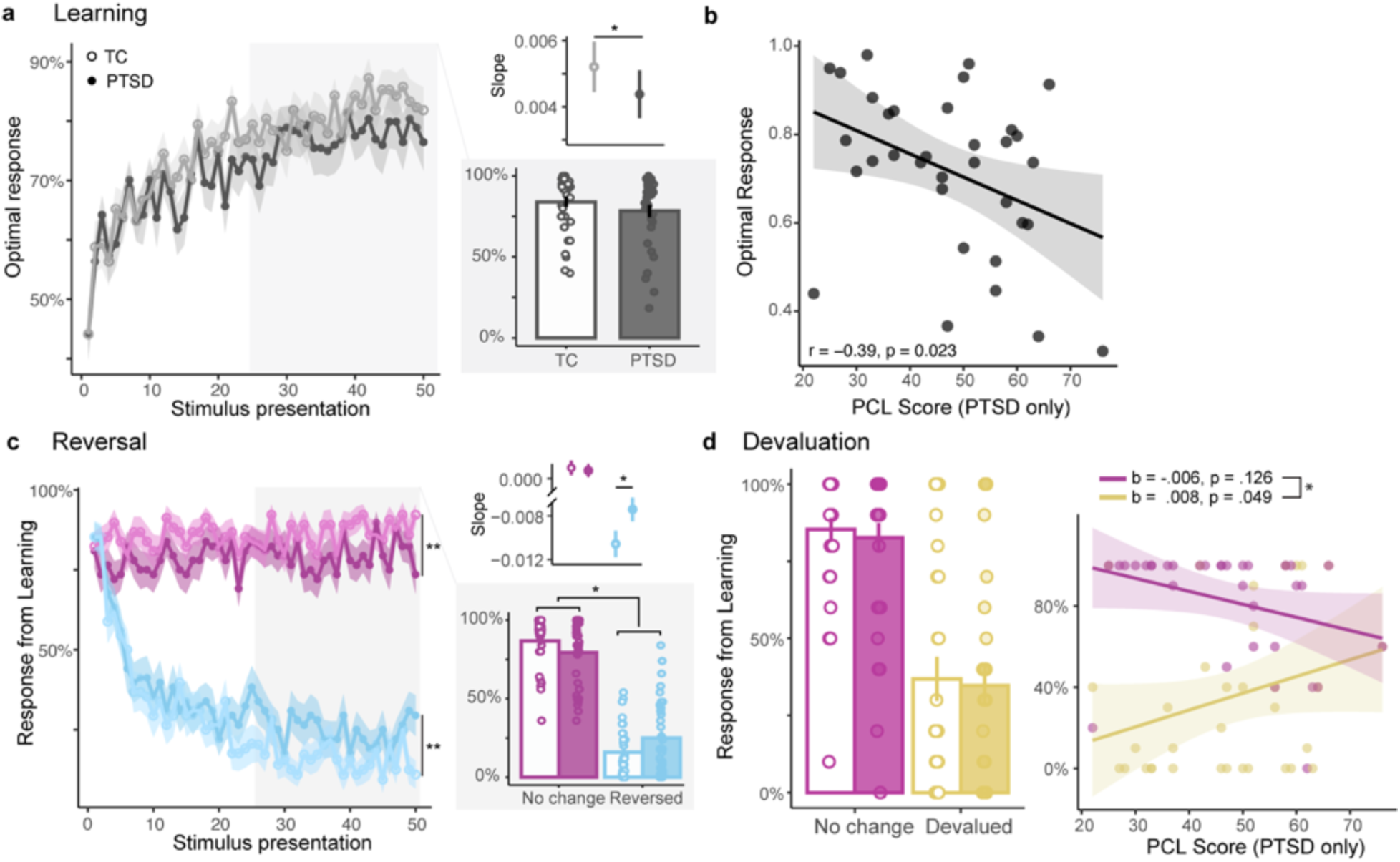
Differences in S-R learning and flexible updating associated with PTSD. **a.** Mean optimal responding for TC group (open circles) and PTSD group (closed circles). Insets shows learning slope for each group (top) with TC showing a steeper learning rate than PTSD, and overall performance in the second half of learning (bottom) with comparable behavior between groups. Ribbons and error bars = ± 1 SEM. **b**. More severe PTSD symptoms are associated with worse performance during learning. Shading = 95% CI. **c.** Maintained responding from Learning phase into Reversal phase for un-changed (purple) and reversed (blue) stimuli. Insets show performance slopes (top), with significantly steeper slopes for TC for reversed stimuli, and overall performance during the second half of learning (bottom) with groups showing opposite patterns for unchanged and reversed stimuli. Ribbons and error bars = ± 1 SEM. **d.** Maintained responding from Learning into Devaluation. No significant difference between groups for unchanged or devalued stimuli. Error bars = ± 1 SEM. **f.** More severe PTSD symptoms are associated with more perseveration for devalued (yellow) but not unchanged (purple) stimuli. Shading = 95% CI. \**p* < 0.05.

To determine whether groups differed in learning trajectories, we used a linear mixed effects model to explain optimal responses over the course of Learning as a function of stimulus presentation and group with subject as a random effect. As anticipated, participants were more likely to make the optimal response as the number of stimulus presentations increased (main effect stimulus presentation: *F*(1,3330) = 558.84, *p* < 0.001) and, as at the end of Learning, there were no overall group differences (main effect group: *F*(1,66) = 0.14, *p* = 0.713). However, groups learned at different rates (group x presentation: *F*(1,3330) = 4.19, *p* = 0.041; **Fig. 2a**). Although both groups improved with practice, the slope of learning was lower in PTSD (TC: *b* = 0.005 [se = .0003]; PTSD: *b* = 0.004 [.0003]; *z*(3330)= −2.05, *p* = 0.041). Effects remained significant when including age as a covariate, and age alone did not significantly predict optimal responses (*F*(1,65) = 3.08, *p* = 0.084).

#### Learning differs with PTSD symptom severity

As learning processes have been shown to differ with PTSD symptom severity ^42^; we examined performance as a function of symptoms in the PTSD group. Individuals with more severe PTSD showed impaired learning (**Fig. 2b**; *r*(33) = −0.389, *p* = 0.023; Supplementary Fig. 5a). Exploratory analyses reveal that this pattern was most strongly related to arousal-related symptoms (PCL subscore for cluster E: *r*(33) = −0.445, *p* < 0.008; Supplementary Fig. 5b). These patterns were not evident in the TC group (PCL score: *r*(33) = 0.016, *p* = 0.930).

### Individuals with PTSD have difficulty flexibly updating learned responses

Our primary hypothesis was that individuals with PTSD would exhibit stronger habit expression and more behavioral rigidity ^12^. Accordingly, we examined how participants adapted to the reversal of a previously learned contingency ^49,52^. Consistent with our hypothesis, individuals with PTSD had more difficulty updating their responses (**Fig. 2c**). Using a linear mixed effects model, we analyzed persistence in previously reinforced responses as a function of group, stimulus type (unchanged or reversed), and stimulus presentation. We found that groups differed significantly in their Reversal trajectories (**Fig. 2c**; group x stimulus presentation: *F*(1,6626) = 3.92, *p* = 0.048) and, crucially, that performance differed by stimulus type (stimulus presentation x stimulus type x group: *F*(1,6626) = 4.02, *p* = 0.045). Compared to the TC group, the PTSD group had greater difficulty updating their responses for the two stimuli for which responses had reversed (TC: *b* = −0.009 [0.0005]; PTSD: *b* = −0.007 [0.0005]; *z*(6626) = 2.82, *p* = 0.005). However, both groups showed similar trajectories for the stimuli for which responses did not reverse (TC: *b* = 0.0007 [0.0005]; PTSD: *b* = 0.0007 [0.0005]; *z*(6626) = −0.012, *p* = 0.991). Effects remained significant when including age as a covariate, and age alone did not significantly predict persistence (F(1,65) = 0.70, p = 0.407).

These data indicate that the PTSD group updated previously learned responses more slowly that the TC group. An analysis of the group by stimulus type interaction (*F*(1,6626) = 7.08, *p* = 0.008) showed that participants in the PTSD group had more persistent responses to the reversed stimuli overall (*z*(66) = 3.06, *p* = 0.002) while showing less persistence for the unchanged stimuli (*z*(66) = −3.30, *p* = 0.001). This suggests that the PTSD group had more difficulty updating previously learned responses when needed, rather than a general tendency to persist in response patterns. This effect lasted even to the end of the Reversal phase (**Fig. 2c**; second half of reversal; stimulus type x group: *F*(1,128) = 5.75, *p* = 0.018). Follow-up analyses further indicated that these impairments were driven by difficulty with flexible updating (Supplementary Note 1).

Finally, we examined whether groups differed in their ability to flexibly respond to changes in outcome value, a classic test of habits ^51^. Contrary to our hypothesis, we did not observe significant differences (*F*(1,128) = 0.178, *p* = 0.673; **Fig. 2d**).

#### Flexible value updating differs with PTSD symptom severity

Reversal behavior was not significantly associated with PTSD symptom severity (optimal response to reversed stimuli: *r*(32) = 0.08, *p* = 0.644). However, more severe PTSD symptoms were associated with greater difficulty in flexibly adapting to devaluation, as evidenced by a greater tendency to make responses that would result in devalued outcomes (PTSD: PCL score x stimulus type: *F*(1,32) = 6.68, *p* = 0.015; PCL score x devalued stimuli only: *b* = 0.008 [0.004], *p* = 0.049; TC: PCL score x stimulus type: *F*(1,28) = 1.20, *p* = 0.277; **Fig. 2d**). Effects remained significant when including age as a covariate, and age alone did not significantly predict devaluation behavior (F(1,31) = 3.19, p = 0.084).

### Putamen engagement tracks learning, flexibility, and PTSD

We hypothesized that the putamen would support gradual acquisition of probabilistic S-R associations, and that this system would be enhanced in PTSD. To test this, we modeled putamen activation per trial as a function of group, Learning run (Early vs Late), and response type (optimal vs nonoptimal) with subject as a random effect. First, we confirmed that the putamen was engaged during learning, finding greater signal on trials for which participants made optimal compared to nonoptimal responses (*F*(1,168) = 14.96, *p* < 0.001; **Fig. 3a**) and a numerically larger increase in engagement from early to late learning (*F*(1,168) = 2.50 *p* = 0.115; **Fig. 3b**). Critically, individuals with PTSD had significantly greater putamen engagement during learning (*F*(1,60) = 6.95, *p* = 0.011). These patterns of putamen engagement were also supported by a whole-brain analysis (Supplementary Fig. 3).

**Figure 3.**
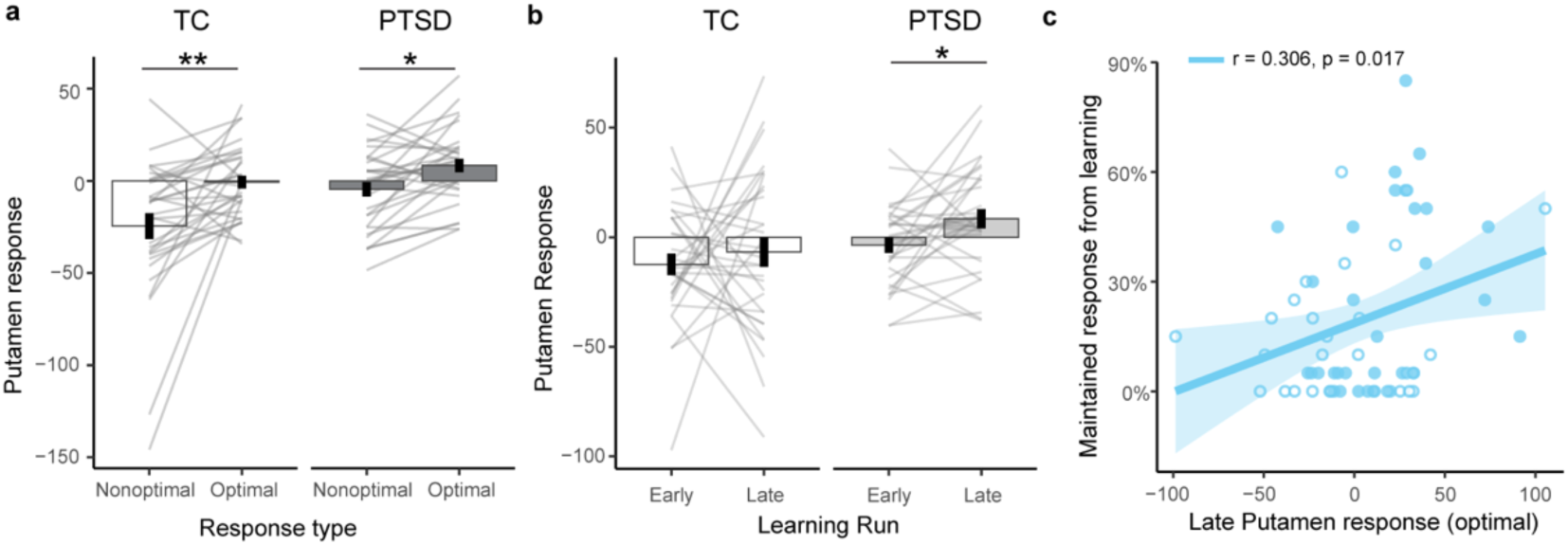
Greater putamen engagement associated with S-R learning, PTSD, and inflexibility. **a.** Putamen responses to optimal and nonoptimal responses during Learning separately for each group; open circles represent TC and close bars represent PTSD. Error bars = ± 1 SEM. Across groups, putamen responses were significantly greater for optimal versus non optimal responses. **b.** Putamen responses in Early and Late Learning. Post-hoc analyses showed significant increases in the PTSD group. **c.** Correlation between putamen responses in Late Learning and later inflexibility (maintained responses in late Reversal). Open circles = TC, filled circles = PTSD. Shading = 95% CI. \*\**p* < 0.01; \*\**p* < 0.05.

We next tested whether putamen engagement during learning had consequences for later flexible updating. We found that greater putamen engagement later in Learning predicted more inflexibility at the end of reversal (**Fig. 3c**; *F*(1,56) = 6.00, *p* = 0.017; also significant when controlling for group differences). We did not find significant associations with behavior following devaluation. These results indicate that the PTSD group engaged the putamen more during S-R learning, and that such reliance was associated with greater difficulty in flexibly changing learned responses.

#### Putamen engagement differs with PTSD symptom severity

Although the relationship with overall symptom severity was not significant (*r*(32) = 0.18, *p* = 0.329), individuals with more severe arousal-related symptoms showed greater putamen engagement by the end of learning (*r*(33) = 0.381, *p* = 0.035; Supplementary Fig. 5b). This association was not evident in the TC group (*r*(33) = 0.041, *p* = 0.827).

### Computational model parameters reveal distinct mechanisms for learning and updating in PTSD

Our results so far reveal that individuals with PTSD were slower to learn probabilistic S-R associations, engaged the putamen more strongly during learning, and had more difficulty updating associations once formed. These findings suggest that individuals with PTSD may engage different underlying strategies when building these associations.

To gain a mechanistic process-level account ^53–55^, we used reinforcement learning models to explain participants’ trial-by-trial responses during learning. We fit these models (described in Supplementary Table 3 and Supplementary Fig. 4) within a hierarchical Bayesian framework which increases sensitivity to individual differences by pooling information across participants within groups ^54,56^.

We found that the same reinforcement-learning model had recoverable parameters and best fit trial-by-trial data for both groups. This model included three free parameters: learning rate (α_+_, with higher levels indicating faster updating of response value after receiving positive feedback for a given stimulus), inverse temperature (β), and trial-level perseveration (*p*; with negative values indicating a tendency to switch responses on consecutive trials; parameter distributions in **Fig. 4a** with details in Methods and Supplementary Table 3). Simulation from posterior parameter estimates successfully recapitulated learning differences (**Fig. 4b**) with strong parameter recovery (TC: all *R*^2^ > .51, all *p*s < 0.001; PTSD: all *R*^2^ > 0.75, all *p*s < 0.001; Supplementary Fig. 4c). A comparison of group-level posterior distributions indicated that the PTSD group had generally lower α_+_ (slower value updating after surprising rewards) and more negative *p* parameter (more erroneous response switching; **Fig. 4a**), though 95% highest density intervals included zero, indicating that group-level differences did not reach significance (Supplementary Fig. 4c).

**Figure 4.**
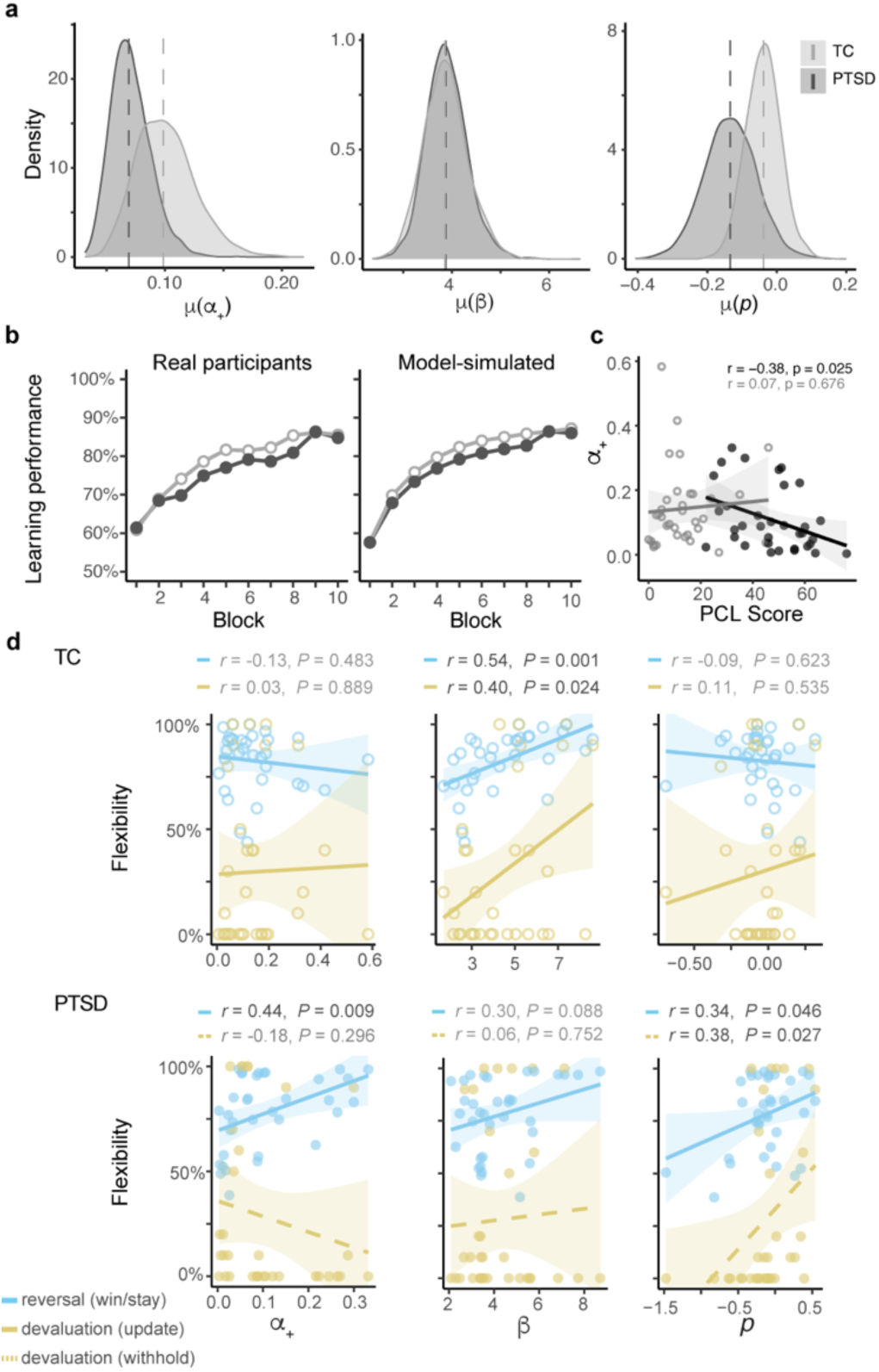
Distinct computations associated with learning and updating in PTSD. **a.** Group-level posterior distributions for model parameters shown separately for TC (light) and PTSD (dark). Dashed lines = posterior means. **b.** Optimal responses over blocks for real (left) and model-simulated (right) data. **c.** Correlations between individual subject level parameters (mean of subject-level posteriors) and flexible behavior in reversal (blue) and devaluation (gold). Shading = 95% CI.

To better understand these parameters, we characterized the behaviors associated with each parameter (Supplementary Fig. 4d). First, as anticipated, *p* strongly correlated with participants’ tendency to switch responses on consecutive trials during learning (TC: *r*(30) = −0.759, *p* < 0.001; PTSD: *r*(30) = −0.916, *p* < 0.001; difference: *z* = 2.20, *p* = 0.028). In PTSD, successful learning was associated with relatively faster value updating (α_+_), but this was not the case for TC (TC: *r*(30) = 0.106, *p* = 0.552; PTSD: *r*(32) = 0.664, *p* = 0.001; difference: *z* = −2.68, *p* = 0.007). In both groups, successful learning was associated with a greater influence of value on choice (higher β; TC: *r*(30) = 0.484, *p* = 0.003; PTSD: *r*(30) = .441, *p* = 0.009; difference: *z* = 0.213, *p* = 0.832).

We reasoned that these parameters, reflecting latent strategies when building probabilistic S-R associations, would have consequences for later flexible updating. Accordingly, we tested how participant-level parameter values from learning predicted later performance. For individuals with PTSD, faster value updating during learning (higher α_+_) and less maladaptive response switching (higher *p*) each predicted more adaptive behavior in reversal (α_+_: *r*(32) = 0.44, *p* = 0.009; *p*: *r*(32) = 0.34, *p* = 0.046; **Fig. 4c**). Higher *p* values again predicted more adaptive behavior after devaluation (response withholding; *r*(30) = .379, *p* = 0.027). These results suggest that, in PTSD, slower value updating and more erroneous trial-level switching predicted more rigid habit-like behavior.

In contrast, for TC, greater influence of value on choice during learning (higher β) predicted more adaptive behavior in reversal (*r*(30) = 0.54, *p* = 0.001) and devaluation (response updating; *r*(30) = 0.40, *p* = 0.024). This suggests that, for TC, value exploitation represented an adaptive strategy for both learning and flexibly updating.

#### Lower learning rate associated with more severe PTSD

As noted earlier, PTSD symptom severity was associated with impaired learning (**Fig. 2b**). Using our computational model, we found that more PTSD symptom severity was particularly associated with slower value updating (**Fig. 4c**; lower α_+_: *r*(33) = −0.384, *p* = 0.025; arousal-related symptoms only: *r*(33) = −0.493, *p* = 0.003; Supplementary Fig. 5a-b). This suggests that individuals with more severe PTSD may rely less on positive feedback to update learned associations.

## Discussion

The potential role of habits in PTSD has been the subject of much theoretical inquiry yet has not been directly tested ^15,16,18–20,33,57^. Here we developed a new behavioral paradigm to test key features of habits – gradual trial-and-error learning of probabilistic stimulus-response (S-R) associations ^13,47^, difficulty remapping responses ^49,50,58^, and difficulty responding to changes in outcome value ^51^ – to quantify potential differences in habit-like processing in PTSD. We found that, although individuals with PTSD were slower to acquire S-R associations, they also demonstrated more rigid habit-like difficulty when needing to change their responses. We further showed greater engagement of the putamen during learning, a pattern that was particularly pronounced in PTSD and was associated with greater subsequent rigidity. Finally, computational modeling revealed that rigidity may arise from differing latent learning strategies in PTSD. These results provide some of the first behavioral, neural, and computational evidence of enhanced habit expression in PTSD.

Despite robust evidence across species that stress facilitates the expression of habits ^23,59,60^, the relationship between stress and the acquisition of habits has been mixed. Some studies suggest that acute stress enhances the rate of S-R acquisition ^34,36,37,61^, although others indicate that acute stress does not influence learning ^24^. Here we found that, although both groups eventually acquired S-R associations, individuals with PTSD learned more slowly than individuals who experienced trauma but did not meet diagnostic criteria (TC group). This pattern of slower S-R learning is consistent with effects of distal stress years or decades prior to the event (for example, early life stress and cumulative life stressors) ^26,27^. Our findings also underscore the dissociation between acquisition and expression of habits. Consistent with this idea, one study found that elevated emotional arousal at encoding both slowed acquisition and enhanced later expression of habit-like responses ^38^.

It is possible that the reward-seeking nature of our task contributed to learning differences in PTSD. We focused on reward learning because habit formation has canonically been studied in reward-based reinforcement learning paradigms, and because reward processing may contribute to PTSD ^62^. However, trauma itself may alter reward processing – for example, rodents exposed to high stress were impaired at transferring positive outcomes from conditioning to later instrumental behavior ^63^. Given emerging evidence for links between altered reward processing and PTSD symptoms ^64–66^ as well as dysfunctional dopaminergic signaling associated with reward learning in PTSD ^64,67,68^, it is possible that alterations in reward processing may have interfered with learning in our task. Thus, if our task were in the aversive domain (i.e., avoiding negative outcomes rather than pursuing rewarding outcomes), individuals with PTSD may have learned more rapidly (but see ^44^). This would be consistent with past findings showing facilitated aversive avoidance learning with prior stress ^34,37,69^, and computational evidence that patients with mood and anxiety disorders learn more slowly from rewards but more rapidly from punishment ^70^. Our findings contribute to a growing literature that considers brain and behavioral biases in PTSD that are more general, that is, not specific to fear-related stimuli^71^. For habits specifically, the neural circuits driving habitual behaviors are distinct from those controlling fear, and thus efforts to reduce such symptoms have been resistant to fear-related interventions such as extinction^5,72^. Further work is needed to characterize avoidance learning in PTSD. Further work is needed to characterize avoidance learning in PTSD.

Consistent with our hypotheses, we find that, despite learning more slowly, individuals in the PTSD group exhibited significantly more perseveration when needing to update previously learned associations. This finding is consistent with recent evidence that acute stress ^28,73–75^ (but see contradictory evidence in rodents ^76,77^) and distal or chronic stress ^28,73^ impairs reversal in both instrumental and classical conditioning paradigms. Successful reversal may serve as a useful read-out of the ability to disengage from ongoing behavior and update actions in service of changing environmental contingencies ^52,78^. When individuals with PTSD continue to engage in rigid safety seeking and avoidance behaviors in response to innocuous stimuli, they may be exhibiting precisely this reversal deficit. Interestingly, ongoing work also suggests that other symptoms such as dissociation may be a form of habitual experiential avoidance^79^.

We also found some support for the idea that PTSD is associated with decreased sensitivity to changes in outcome value. In this canonical test of habits ^51^, we did not uncover group differences, but did observe that individuals with more severe PTSD engaged in more habitual responding. Although stress exposure leads to insensitivity to changes in outcome value in both rodents and humans ^22–25^, recent evidence suggests that gradations of stress severity or appraisal may be impactful. Both higher levels of early life stress ^69,80^ and greater stress-induced negative affect ^81^ were associated with less sensitivity to devaluation. Our finding that more severe PTSD tracked less sensitivity to devaluation aligns with these results and suggests that continuous measures of stress severity may be a relevant predictor for habitual behaviors in devaluation. It is also worth noting that devaluation procedures are tricky in humans ^82,83^ and typically require extensive training. Thus, it is possible that our design did not include sufficient trials to observe a robust group difference. Evidence of group differences in reversal, but not devaluation, is also consistent with recent work showing that response remapping (i.e., reversal) is a more consistent measure of habit emergence than response withholding (typically measured after devaluation) ^50,58,84^. These considerations underscore the importance of including multiple assays, like both reversal and outcome devaluation, to understand habit-like rigidity.

The dorsolateral striatum (human putamen) plays a crucial role in habit formation and expression ^13,85–87^. Engagement of this structure has been shown to be particularly pronounced later in S-R learning, corresponding to stronger and more habit-like associations ^12,14,88,89^. Consistent with these findings, we showed greater putamen engagement when responding optimally in our S-R task. Importantly, the dorsolateral striatum has been implicated in the effects of stress on habitual behaviors; stress alters fronto-striatal reorganization, heightens amygdala-striatal connectivity, enhances neural representations for motor response over goals, and increases dendritic complexity and arborization in the dorsolateral striatum ^20,22,29–31^. Such changes may be particularly important in PTSD. One study found that altered cortical-striatal connectivity was associated with PTSD symptoms, but not trauma exposure alone ^32^. Here we show greater putamen engagement in PTSD relative to trauma exposure alone as individuals build habit-like S-R associations. As the putamen is associated with gradual trial-and-error based learning ^86,90^, the relatively slower strategy of learning and updating from positive feedback in PTSD may have contributed to enhanced engagement of this structure when responding correctly. Greater putamen engagement in PTSD is also consistent with theories of PTSD positing that enhanced dorsolateral striatum engagement may underly lasting maladaptive behaviors in response to trauma-related cues ^19,33,57^. Although more work is needed to understand the cellular and signaling mechanisms that give rise to enhanced putamen engagement in PTSD, our work argues for more consideration of the striatum and associated behaviors in theoretical models of PTSD.

In addition to differing choice tendencies, PTSD has been theorized to involve distinct latent processes that affect how these individuals learn and represent information ^42,44,71,91^. Several studies have employed computational modeling approaches, including reinforcement learning models, to characterize learning strategies in PTSD ^42–44,92^. Our study expands on these findings by providing evidence for dissociable relationships between learning strategies and subsequent habit-like rigidity in PTSD. During learning, both groups had comparable inverse temperature parameters (perhaps reflecting that the value-updating algorithms employed in the model were well-suited to choices in both groups ^93^). Notably, higher inverse temperature predicted subsequent behavioral flexibility in the TC group, suggesting that these stable value-based choice policies were adaptive; thus, these individuals may have relied on a more exploitative policy to guide subsequent response remapping. Although other parameter differences were not statistically significantly different during learning, group-level estimates for learning rate were lower and trial-to-trial switching tendencies were more extreme in the PTSD group. As noted above, lower learning rate may reflect a greater reliance on gradual trial-and-error updating characteristic of the dorsolateral striatum ^90,94^. Interestingly, individuals within the PTSD group who were able to achieve higher learning rates were also more able to flexibly update behaviors. This raises the possibility that accommodating these learning rate differences, perhaps by extending therapy duration, may be particularly beneficial for individuals with PTSD. Finally, regarding trial-to-trial response switching, this inappropriate use of responses to one stimulus to inform responses to another independent stimulus may reflect a lack of sensitivity to state changes (see also ^92^). This may be consistent with a tendency to respond inappropriately in non-trauma-related contexts. Indeed, more extreme values of this switching parameter were associated with greater difficulty in flexibly updating responses later. Together, these results are consistent with previous work demonstrating that individuals with PTSD leverage different latent learning strategies, with important differences in the process of learning as opposed to impairments in learning per se. Although further work is needed to examine these computations in other learning domains, our findings suggest that gradual learning and difficulty discriminating contexts may play important roles in the development of habits in PTSD.

Our results suggest that individuals with PTSD employ different latent learning strategies that manifest as slower learning, greater engagement of the dorsolateral striatum, and later expression of rigid habit-like behaviors. It should be noted that our sample was predominantly female. As the majority of PTSD research has been conducted in males, this work makes an important contribution by examining females, who also exhibit higher lifetime prevalence rates ^95^. Nevertheless, this sex imbalance may limit generalizability. Together, our findings offer insight into the mechanisms by which habits are expressed in PTSD, which may explain how certain pathological behaviors are acquired and expressed. Habitual control of behavior, and its consequences for the ability to flexibly update actions, may reflect an important dimension of PTSD and serve as a clinically meaningful target.

## Methods

### Sample characteristics

A total of 70 participants (N = 35 meeting diagnostic criteria for PTSD and N = 35 reporting trauma exposure but not meeting criteria, details below) recruited from the New Haven community completed study procedures. These group sizes were determined based on a power analysis showing significant stimulus-response learning in an established probabilistic task (80% power, alpha = 0.05) ^21,45^. All participants were fluent in English, had normal or corrected to normal vision, and provided written informed consent. Exclusion criteria included: moderate/severe traumatic brain injury, bipolar disorder, learning disability, neurological disorders, use of antipsychotic, hypnotic, sedative, or anti-addiction medication, current severe substance use disorder, and metal in body (for MRI safety). Diagnostic criteria for PTSD were assessed using the Structured Clinical Interview for DSM-5 (SCID-5) administered by a trained interviewer (DTN). All study procedures were approved by the Yale University Institutional Review Board.

### Task design

To determine whether individuals with PTSD differed in habit-like learning, we designed and implemented a multi-phase probabilistic reinforcement learning task consisting of 3 distinct phases: (a) Learning; (b) Reversal; and (c) Devaluation (**Fig. 1**). Participants completed the task while in the MRI scanner, with the images viewed via a mirror attached to the head coil and responses made using an MRI-compatible button box. In each phase, participants were instructed to place a single visually presented stimulus object (e.g. stapler) into the correct room (Left or Right door) by making two button presses. Participants had less than two seconds to first select a door (first button press), and then less than half a second to open it (second button press). Each stimulus object was presented in the center/top of the screen, and the same two door options were presented on all trials (**Fig. 1a**). Choosing the correct door for a given stimulus resulted in presentation of a fruit image that signaled earning 10 points (which were added to their monetary bonus). Fruits were stimulus-specific; that is, the correct responses for a given object would always earn the same fruit. If participants did not complete the first and second button presses to select and open the door within the time limit, this resulted in a loss of 1 point. The length of the inter-trial intervals was jittered such that all trials were the same length.

To target striatal-mediated trial-and-error learning, the stimulus-room associations were probabilistic with a reinforcement rate of 80% ^45,47^. Participant learned 6 stimulus-room associations with each room designated as the optimal response for 50% of the objects. This number of associations was chosen to minimize reliance on working memory and to allow subsets of associations to change in subsequent phases ^48^. Trials were pseudorandomly interleaved such that the same object could not be presented on more than 2 consecutive trials, and all stimuli were presented 10 times per block. Invalid (20%) feedback did not occur first in any block or immediately following valid (80%) feedback for the same stimulus.

The Learning phase consisted of 300 trials over 10 blocks, with 5 blocks per scan run. Immediately thereafter, the Reversal phase began, and two of the six objects switched their room association (i.e., Right door → Left door for one stimulus and Left door → Right door for another stimulus). This change in stimulus-room associations was not signaled to the participant. The Reversal phase also contained 300 trials over 10 blocks. Finally, participants underwent a Devaluation procedure, in which they were which two of the six fruits were now “rotten,” and that they needed to avoid earning them. The two reversed objects were never devalued. If participants made a response to an object that would result in one of these fruits, there would be a 10-point deduction from their score (and thus a decrease to their monetary bonus). After these instructions, participants resumed placing objects in different rooms without receiving feedback. Participants first completed 6 trials to practice responding without feedback, followed by 1 block (30 trials) assessing the impact of outcome devaluation.

To confirm that participants perceived the fruit outcomes as valuable, and subsequently adjusted value for fruits that were rotten in devaluation, participants completed five brief consumption tests preceding every functional imaging run (Supplementary Fig. 1a) ^96,97^. Here, participants were instructed to select fruit to add to their score. All stimulus-unique fruit (6 total) were presented three times each, sequentially, and all in a random sequence, with participants having under 500ms to respond (via a single button press) to each fruit. A yellow circle appeared around fruits that participants selected.

To confirm that participants learned stimulus-outcome associations, at the end of the experiment participants completed a contingency awareness test. Participants were instructed to select the stimulus-unique fruit that could be earned for each object stimulus. Each of the six object stimuli were presented for three seconds. For each object stimulus, participants were shown each of the six fruit stimuli that could have been paired with that object stimulus in sequence and had two seconds for each fruit stimulus to select it as a match (Supplementary Fig. 1b).

Two participants were excluded from all behavioral analyses, one whose behavioral data was missing (TC), and another whose behavioral data were incomplete (PTSD), both due to technical errors. Two further participants (both TC) were excluded from analyses of behavior after the Learning phase, one due to a technical error involving task administration, the other due to not completing the task.

### Experimental procedures

Prospective participants from the greater New Haven community were first phone-screened to determine eligibility for participating in the experiment. The study then included 2 sessions on two separate days, including an intake session and a scanning session, no more than 163 days apart. At intake, participants completed self-report measures (PTSD Checklist for DSM-5 to assess symptom severity, and the Life Events Checklist for DSM-5 to assess history of traumatic events) and a clinical interview with a trained interviewer to determine if they had experienced a Criterion A traumatic event and/or met diagnostic criteria for PTSD.

Eligible participants then came in for a second session on a separate day to complete the probabilistic learning task while undergoing MRI. Participants were asked not to consume alcohol for 24 hours prior to the MRI session. Participants first completed a training session with the experimenter (DTN or KBL) outside of the MRI. During this training session, they were instructed to learn where objects belong (left or right door), were made aware that reinforcement was probabilistic, and that stimulus-response associations could reverse at some point during the task. They also had the opportunity to ask questions practice making multi-button responses to individual objects. Subjects practiced learning 3 associations that included probabilistic feedback and a reversal; practice continued until participants demonstrated successful performance. Finally, participants completed a post-practice comprehension quiz to ensure that all facets of the task were understood. Participants could not advance to the scan without passing this quiz.

### Magnetic Resonance Imaging

***Acquisition parameters.*** Functional MRI (fMRI) data were collected while participants completed the MPL task. MRI data were acquired using one of two Siemens 3 Tesla Prisma scanner using a 64-channel head coil at the Yale Magnetic Resonance Research Center at Yale University. We confirmed that groups did not differ significantly in scanner assignment (χ^2^(2) = 1.14, p = 0.565).

We acquired a high-resolution 3D T1-weighted anatomical MPRAGE image (TR = 2,400ms, TE = 1.22ms, voxel size = 1 mm^3^, flip angle = 8°, FOV = 256×256, 208 slices). Functional data were collected using an echoplanar imaging (EPI) sequence (TR = 1,000ms, TE = 30ms, voxel size = 2mm^3^, flip angle = 55°, multi-band factor = 5, FOV = 220×220).

***Preprocessing.*** fMRI data were minimally preprocessed using FSL (6.0.3). Data were first skull-stripped using the Brain Extraction Tool (BET), pre-whitened using FMRIB’s Improved Linear Model (FILM), motion corrected (MCFLIRT) and high-pass filtered at 0.01Hz to remove low-frequency signal drift. We then aligned functional data to a reference functional image, and to standard (MNI) space using boundary-based registration. Six rigid-body motion parameters were derived using FSL Motion Correction using FMRIB’s Linear Registration Tool; MCFLIRT.

Eight participants were excluded entirely from neuroimaging analyses: incomplete or absent behavioral data (see above; TC: N = 1; PTSD: N = 1), missing T1 structural scans due to early termination of the scan session (TC: N = 1; PTSD: N = 1), missing functional imaging data due to discomfort in scanner (TC: N = 1; PTSD: N = 1), and excessive head motion during all learn runs (mean framewise displacement > 1.5mm; TC: N = 1; PTSD: N = 1). A further early learning run was excluded from one PTSD participant due to excessive head motion.

### Analysis: Behavior

Behavioral analyses were conducted in R (version 4.2.1). Linear mixed effects models were computed using the lme4 (1.1.30) ^98^ and lmerTest package (version 3.1.3) ^99^ with post-hoc analyses using emmeans (version 1.8.2) ^100^. Early and Late epochs for the Learning and Reversal phases were defined by fMRI run, with each run comprising 150 trials.

***Task validation***. To validate acquisition for both groups during the Learning and Reversal phases, we computed mean optimal response rates per participant across stimulus types (unchanged, to-be reversed, to-be devalued), within early and late runs per phase. We used one-sample t-tests to determine whether optimal responses differed significantly from chance (50%). P-values were corrected for multiple comparisons using the Benjamini-Hochberg FDR procedure. To validate devaluation, we first tested whether participants in each group successfully decreased the value of the “rotten fruits” using a paired samples t-test comparing the proportion of valued versus devalued fruits that each participant selected in the post-devaluation consumption task (Supplementary Fig. 1a). To test whether participants successfully updated responses to the stimuli associated with devalued outcomes, we ran a paired samples t-test comparing the proportion of original responses for devalued versus unchanged stimuli.

Lastly, we investigated whether groups differed in their memory of stimulus-outcome associations using a linear mixed effects model, predicting accuracy (correct stimulus-outcome match) as a function of stimulus type, group, and their interaction. Post-hoc one sample t-tests against chance were used to verify that accuracy was above chance for each stimulus type per group.

***Learning***. We analyzed optimal responses as a function of stimulus presentation (repetitions of each object stimulus), group (TC vs PTSD), and their interaction, with subject as a random effect. Significant interactions were further examined using emtrends. To compare final achieved performance between groups, we ran an independent samples t-test comparing optimal responses in Late learning. We also computed the number of stimulus-response associations [0-6] learned per subject by running a binomial test comparing optimal response rate to chance (50%) for each stimulus (threshold: p < 0.05). The number of correctly learned stimuli was then entered into a linear model with group as a predictor. Lastly, we tested if optimal responding was associated with symptom severity (overall PCL score) using an ordinary least squares (OLS) linear regression model, separately for each group. Exploratory analyses of associations between learning behavior and PCL subscores (e.g. arousal) were performed using the same method.

***Reversal***. To investigate how well participants flexibly updated previously acquired responses, we calculated the proportion of responses that participants maintained – rather than updated – from Learning into Reversal. We then ran an LME predicting maintained responses during Reversal based on stimulus presentation, stimulus type (unchanged, reversed), group, and their interactions with subject as a random intercept. We then compared final achieved performance in Late Reversal between groups using an independent samples t-test.

***Devaluation.*** We calculated the rate of maintaining (rather than updating) originally acquired responses for each stimulus type during devaluation. Maintained responses were analyzed as a function of stimulus type (unchanged, devalued), group, and their interaction with subject as a random intercept. We used an OLS linear regression to predict original responses for the unchanged and devalued stimuli separately as a function of symptom severity.

### Analysis: fMRI

To investigate the involvement of the dorsolateral striatum in each group during Learning, we ran first-level models estimating evoked responses to choices in FSL (FEAT) and group-level models in AFNI (23.1.07, 3dLMEr).

Preprocessed functional runs in native space were entered into a general linear model (FEAT) with 3 main regressors: optimal response trials, non-optimal response trials, and skipped trials as well as the six rigid-body motion parameters as control regressors. Each trial was modeled as a boxcar beginning at stimulus onset and ending at choice onset (1-2 TRs, based on RT) convolved with a double-gamma hemodynamic response function. For these analyses, 10 participants (TC: N = 6; PTSD: N = 4) had insufficient nonoptimal trials in the second learning run (< 3 trials) and were excluded.

***Region of interest (ROI) analysis.*** To test the role of the putamen, we defined a participant-specific anatomical putamen ROI using each participant’s MPRAGE (FSL’s FIRST segmentation). We then extracted mean output beta maps (see above) from this ROI for optimal and non-optimal responses from early and late learning. These putamen beta maps were then modeled as a function of run (early learning, late learning), response type (optimal, non-optimal), and group (TC, PTSD), with subject as a random intercept.

We then assessed whether putamen responses during learning were associated with later evidence for habitual responding. To do so, we focused on late learning, once associations were fully formed (and when putamen has previously been shown to be maximally engaged ^12^). We then used a linear model to test whether stronger putamen responses in late learning predicted maintained responses to changed cues in late reversal and devaluation while controlling for group differences. Finally, we tested if putamen responses in either early or late learning were associated with symptom severity (overall PCL score) using Pearson correlations, separately for each group. Exploratory analyses of associations with PCL subscores were performed using the same method.

***Whole-brain analysis.*** We further validated the role of the putamen using whole-brain analyses. Output beta maps (parameter estimate images) for optimal and non-optimal responses from the first-level models described above were aligned to MNI152 standard space using FLIRT, then smoothed with a 6mm FWHM Gaussian kernel. These beta estimates were then entered into a linear mixed-effects model (AFNI’s 3dLMEr) with neural responses modeled as a function of run (early learning, late learning), response type (optimal, non-optimal), group (TC, PTSD), and their interactions, with subject as a random intercept. Voxels showing main effects of group, response type, and group (late learning only) were identified. These significant voxels from post hoc contrasts were determined using voxel wise FDR-correction at *p* < 0.05.

### Computational modeling

To investigate latent learning strategies, we implemented computational reinforcement learning models to describe trial-level choice (Left, Right) data from the Learning phase (actions were coded as Left = 1, Right = 2). We tested eight models. Of these, seven models were modified variants of the Rescorla-Wagner (RW) model involving error-driven predictive learning and a softmax choice policy. One (null) model was a deterministic win-stay lose-shift rule with no error-driven learning. For all models, skipped trials were excluded from the fitting process.

For model estimation, we used hierarchical Bayesian estimation, implemented in the probabilistic programming language Stan (2.21.7) which uses a Markov Chain Monte Carlo sampling algorithm (HMCMC). Models were fit separately for each group using expected log predictive density (elpd), an estimate of out-of-sample predictive fit via leave-one-out cross validation. Weakly informative priors were used for group-level means and standard deviations (Normal(0,1)). A total of 2,000 samples were drawn following 1,000 warmup iterations for each of 4 chains.

We first implemented a probabilistic win-stay/lose-shift model with inverse temperature to capture choice stochasticity, but no error-driven learning updates. This model captures a simple outcome-driven strategy, where a future action for a given stimulus (stimulus presentation) is based on the most recent outcome for that same action (stimulus presentation – 1). This model served as a non-learning baseline to compare against error driven learning models. First, a desired action was computed according to the equation:

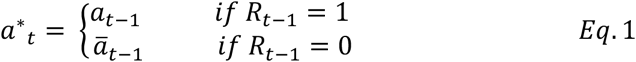

Where *a*^∗^ is the desired action (e.g. Left), and *a*) is the alternative action (e.g. Right). These values were combined into a vector:

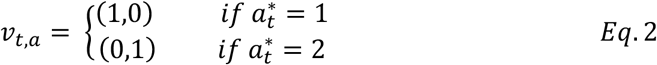

Choices were then generated using a softmax function to convert desired actions into choice probabilities. An inverse temperature (β) adds stochasticity to the choice policy, controlling the extent to which actions follow a win-stay/lose-shift policy.

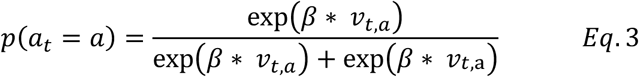

We next implemented a series of error-driven learning models that use Rescorla-Wagner updating rules to update value trial by trial. For these seven models, Values (*Q*) for each of the two actions (Left, Right) are updated separately for each of the six unique stimulus-response associations. All *Q*-values were initialized to 0. *Q*-values for the unchosen door and for non-observed stimuli were not updated. The RW model free parameters differed in (1) learning rates (single learning rate, separate win-learning and loss-learning rates) (2) inclusion/exclusion of trial-level perseveration, and (3) whether value updating was implemented at the stimulus level (adaptive learning) or at the trial level (maladaptive learning).

We first implemented a baseline maladaptive learning model that learns over trials, instead of stimulus presentations; that is, values are updated on consecutive trials, irrespective of the stimulus presented. This trial-level learning model updated values for each door as follows:

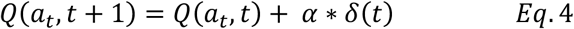

where *α* reflects the strength of prediction error updates to the door values, or the learning rate.

*δ*(*t*) represents the difference between actual and anticipated value computed as:

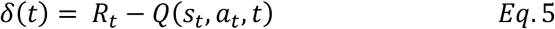

Like the win-stay/lose-shift model, *Q* values were entered into a softmax equation with an inverse temperature parameter to compute action probabilities (see Eq 3.).

We then computed a baseline adaptive learning model, where values are updated over stimulus presentations (instead of trials). Here, values are computed similarly to the trial-level learning (Eq. 4) model, but contingent on the current stimulus *s*. Values were then fed into a softmax equation (Eq. 3).

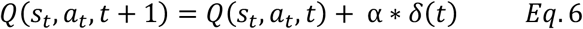

We then wanted to test the hypothesis that individuals exhibit both adaptive (stimulus-level) and maladaptive (trial-level) learning. To do so, we implemented a hybrid learning model: here, separate values were computed over trials and stimulus presentations with a trial-level learning rate, and stimulus-presentation learning rate respectively (see Eq. 4 & 5). Then, a combined door value was computed each trial as a weighted sum of these separate values using a free parameter *w*:

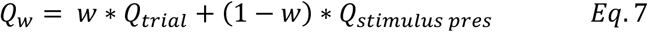

These weighted Q values were then entered into the softmax equation (Eq 3).

Given previous modeling work in humans that has observed distinct updating for rewarded versus non-rewarded trials, and specifically of several observations of outcome-related encoding differences in PTSD ^42,44,65^ we tested whether learning differed for trials that resulted in positive or negative feedback. To do so, we implemented a model with separate learning rates for rewarded trials (a_+_) and non-rewarded trials (a_−_). Value updates for this two-learning rate model were computed using update rules according to Eq. 6 and entered into the softmax equation (Eq. 3).

Although trial-level learning is maladaptive, and did not describe the data well, participants may still exhibit trial-level behavior tendencies that are independent of value updating (i.e. choice autocorrelation such as switching and perseverating ^54,101^. To test this, we included a new free parameter p that scales the value of only the previous choice where positive values indicate perseverating, and negative values indicate switching. We included this parameter *p* in the softmax equation (updated from Eq. 3). Value updates for this model were identical to the above two-learning rate model. Choice probabilities were computed as below:

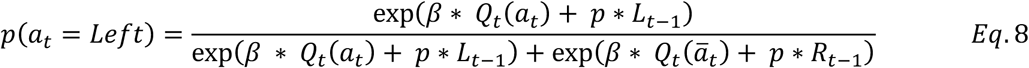

Where *L_t_*_–1_ and *R_t_*_–1_are binary indicator variables equal to 1 if the previous choice was Left or Right, respectively. In the above two-learning rate model, population posterior distributions of a_−_ were very near 0, suggesting that learning rates over non-rewarded trials did not contribute significantly to behavior; moreover, parameter recovery for this model was poor.

To address this issue, we implemented a model with single learning rate for rewarded choices (α_+_), an inverse temperature parameter to capture stochasticity in choice (β), and a perseveration parameter (*p*). In this model, values (*Q*) for each of the two actions (Left, Right) are updated separately for each of the six unique stimulus-response associations. *Q*-values for the unchosen door and for non-observed stimuli were not updated. *Q*-values for both doors on negative feedback trials were not updated. All *Q*-values were initialized to 0. On trials in which the outcome was positive (e.g., fruit presented), *Q*-values were updated according to the following rule:

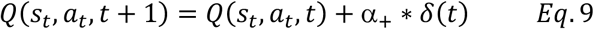

where the prediction error *δ*(*t*) is computed the same as in Eq. 5. Here, *s* is the stimulus (e.g. stapler), *a* is the action (Left, Right), and *t* is stimulus presentation. *R_t_* represents the reward on a given trial (1 if fruit is presented; 0 in the absence of fruit). The learning rate is a constant, free parameter. No update was applied to incorrect (loss) trials; thus, learning was driven only by positive outcome trials. Choice probabilities were generated using a softmax equation (Eq 8), where *p* represents trial-level perseveration (positive = repeat action from previous trial, negative = switch action from previous trial), β reflects the degree that choices track relative Q-values (higher = more exploitative), and *a’* represents the alternate (unchosen) door.

Individual-level parameters were drawn from group-level normal distributions, and later constrained as follows: α_+_: [0,1] via phi approximation (Phi_approx function), β: positive only (exponential transformation) *p*: unconstrained.

The hBayesDM (version 1.2.1) package was used for model diagnostics and comparison. elpd differences (relative to the best-fitting model) are reported for each group (Supplementary Table 3). For this model, visual inspection of MCMC chains confirmed mixing of MCMC samples for each group (Supplementary Fig. 4a). Model convergence was assessed for each group using the R-hat statistic that compares within and between-chain variances, where R-hat values closer to one indicate good convergence. All parameters in both groups yielded R-hat values below 1.05. Posterior distributions of group-level parameter estimates are reported with 95% highest density interval (Supplementary Fig. 4c), computed using the HDinterval package.

***Parameter recovery***. To further validate our model and test identifiability of model parameters, we simulated synthetic behavioral data using participants’ posterior mean estimates and re-fit our winning model using the same approach described above to simulated data. We then compared these recovered values to the true, generating parameters. Our results confirm that parameters in the winning model were recoverable (all *R*^2^ > .506, all *p*s < .001; Supplementary Fig. 4b)

***Posterior predictive checks***. To test the ability of the model to capture group differences in learning, we tested whether behavior generated by the model could reproduce our observed learning trajectories. For each participant, trial-level choices were generated from their posterior predictive distribution (y predicted) and averaged across draws to derive predicted choice probabilities. Optimal choices were derived for each trial for both the simulated, as well as real empirical data, and averaged into pseudo-blocks of 30 trials (due to exclusion of skipped trials in modeling). Visual inspection confirms that simulated behavior was very similar to true group differences in learning trajectories (Fig. 4b).

***Parameter – behavior relationships***. To examine relationships between the derived model parameters and raw behavior, we computed Pearson correlations between individual-subject level parameter estimates (posterior means) and summary behavioral statistics, separately for each group, for each phase (Supplementary Fig. 4d). To confirm that parameter values were sensitive to expected learning behaviors, we quantified the proportion of optimal responses (choosing 80% reinforced option), proportion of win-stay behaviors (trials for which responses following positive feedback were repeated for a given stimulus), and trial-level switching (choosing the opposite door of the previous trial) during learning.

To test whether variability in specific latent learning processes predict later habitual behavior, we assessed the relationship between learning parameter values and behavior during the Reversal and Devaluation phases using ordinary least squares linear regressions within each group, for each phase. In Reversal, we calculated successful flexibility as win-stay rates for reversed stimuli. In Devaluation, we calculated successful flexibility as either withholding (i.e., trials presenting the devalued stimulus on which the participants did not select a door) or updating responses (e.g. selecting Left door to the devalued stimulus if Right door was optimal in Learning) to stimuli for which the outcome was devalued. We highlight key relationships between parameters and flexible behavior for respective conditions (win-stay behavior in Reversal, and updated or withholding response behavior in Devaluation) in Fig. 4c.

## Acknowledgments

Conceptualization: E.V.G. and J.H.K.; Methodology: E.V.G., S.K., and S.M.G.; Investigation: K.B.L. and D.T.N.; Formal analysis: K.B.L. and E.V.G.; Resources: K.B.L. and S.M.G.; Data curation: K.B.L. and D.T.N.; Writing – Original Draft: K.B.L.; Writing – Review & Editing: D.T.N., S.K., J.H.K., S.M.G., and E.V.G.; Supervision: E.V.G.; Funding acquisition: E.V.G. Software: K.B.L., S.M.G., S.K., and E.V.G. The authors are grateful to Erin O’Brien for helpful discussions and the technicians at the Yale Magnetic Resonance Research Center for assistance with MRI data collection. This work was supported by a NARSAD Award from the Brain and Behavior Research Foundation (EVG) and the National Center for PTSD.

## Notes

### Competing Interest Statement

The authors have declared no competing interest.

