## Supplemental Data for "Striatal habit system drives behavioral rigidity in PTSD"

**Supplementary Table 1**

| **Measure** | **Trauma-exposed control  (N = 35)** | **PTSD**  **(N = 35)** | **Difference?^1^** |
| --- | --- | --- | --- |
| Age: M (SD) | 29.6 (7.82) | 31.9 (10.1) | p = 0.28 |
| Sex | M: 34.4% F: 65.7% | M: 25.7% F: 71.4% | p = 0.60 |
| Post-traumatic Stress Disorder  Checklist (PCL-5) | 13.3 (11.2) | 47.4 (13.4) | p < 0.001 |
| Life events checklist (LEC-5) | 3.5 (2.1) | 4.9 (2.4) | p = 0.010 |
| Antidepressant use | 11.4% | 42.9% | p = 0.002 |
| Beta blocker use | 2.9% | 0% | p = 0.221 |
| Corticosteroid use | 0% | 2.9% | p = 1.00 |
| Alcohol use disorders identification test (AUDIT) | 3.9 (4.3) | 7.2 (7.2) | p = 0.026 |
| Endorsed ever experiencing obsessions | 8.8% | 17.1% | p = 0.446 |
| Endorsed ever experiencing compulsions | 8.8% | 25.7% | p = 0.100 |

^1^Demographics compared using t-tests, F-tests, or Chi-squared tests as appropriate. LEC scores refer to total events endorsed as “happened to me”.

**Supplementary Table 2**

|  | **Free parameters** | **TC: ELPD** | **PTSD: ELPD** |
| --- | --- | --- | --- |
| Stimulus-level learning (from positive feedback only) and trial-level perseveration | α_+_, β, *p* | **0.0** | -2.3 |
| Stimulus-level learning (from positive and negative feedback separately) and trial-level perseveration | α_+_, α_-_, β, *p* | -4.5 | **0.0*** |
| Stimulus-level learning (from positive and negative feedback separately) | α_+_, α_-_,β | -25.0 | -127.5 |
| Stimulus-level learning and trial-level learning | α_state_, α_trial,_ β | -261.5 | -240.0 |
| Stimulus-level learning | α_state,_ β | -330.6 | -330.3 |
| Win-stay/lose-shift (deterministic choice policy based on previous trial) | β | -2780.4 | -2588.3 |
| Trial-level learning | α_trial_, β | -2797.7 | -2611.3 |

ELPD = estimated log predictive density. Values are shown relative to best-performing model (assigned score of 0.0), with more negative values indicating worse fits. Best performing model highlighted in bold. *one or more parameters unrecoverable

**Supplementary Note 1: Alternative explanations of reversal impairment in PTSD**

We considered several possible explanations for reversal impairments in PTSD. First, incongruous feedback (to reversed stimuli) following an un-signaled reversal could have destabilized all previously learned responses. However, when comparing performance between late Learning and early Reversal, we saw significantly improved performance for the unchanged stimuli (*F*(1,6730) = 13.04, *p* < 0.001), and this effect did not differ between groups (group x phase: *F*(1,6730) = 0.096, *p* = 0.757; Supplementary **Fig. 2b**). Second, deficits in reversal could simply be explained as impairments in learning as opposed to difficulty with flexibly updating. If so, we would expect reversal to reflect “re-learning”, with groups reaching similar levels of performance by the end of Reversal as they did at the end of Learning. However, if reversal impairments are better explained by a distinct process of inflexible updating, we would expect performance to be different. To test this, we used a linear mixed model predicting optimal responses in the second half each phase by group, phase, and stimulus presentation. Consistent with an impairment in flexible updating, both groups performed differently on the same stimuli from the end of learning to the end of reversal (group x phase: *F*(1,3276) = 24.96, *p* = .045). The PTSD group never regained the same level of performance they had shown at the end of learning when they were required to update their responses (End Learning - End Reversal: *b* = .069 [.013], *p* < .001; Supplementary Fig. 2c). In contrast, the TC group exceeded their prior Learning phase performance, indicating flexible and successful updating (End Learning - End Reversal: *b* = -.027 [.014], *p* = .048). Critically, this dissociation was unique to the reversed stimuli. These results indicate that the observed reversal impairments in PTSD were likely related to difficulty in flexibly updating associations.

**Supplementary Figure 1. Task Comprehension checks.** To validate the participants understood to avoid rotten fruits (negative points), we implemented a consumption task where participants freely select consecutively presented fruits. At the end of the experiment, they also completed a short task to assess memory for stimulus-outcome associations **a.** Consumption Tests: Valued and Devalued fruit selection at each of 5 consumption checkpoints. Following rotten fruit instructions at checkpoint 5, Participants in both groups exhibited successful devaluation of the rotten fruits. **b.** Contingency Test: Stimulus-outcome memory performance. Both groups exhibited above chance memory for stimulus-outcome associations for each of the stimulus types learned throughout the experiment (all ps < 0.001). Groups did not differ in stimulus-outcome memory (group x stimulus type: p = 0.180). Error bars = ± 1 SEM.

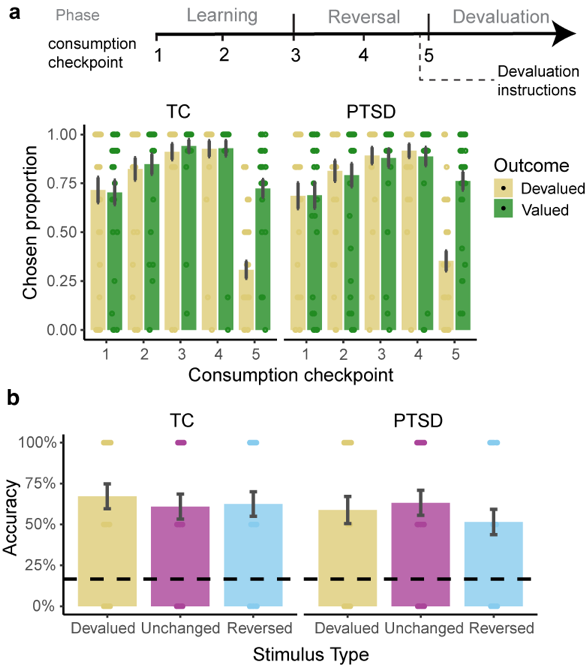

**Supplementary Figure 2. Stimulus learning across phases. a.** Average number of learned stimuli. Groups did not differ in the average number of learned stimuli (between 0 and 6 total per subject) in either of the phases. **b.** Mean optimal responding with standard error for the unchanged stimuli for each group across Late learning and Early reversal. Groups did not differ in performance across phases, showing numerically increased performance from learning to reversal. **c.** Mean optimal responses with standard error as ribbons for the reversed stimuli between the end of learning, and the end of reversal. The TC group demonstrated significant improvement, indicating successful re-learning of reversed stimuli; the PTSD group exhibited significantly worse performance between phases, indicating a failure to successfully update; this effect was specific to the reversed stimuli only. Error bars/ribbons = ± 1 SEM.

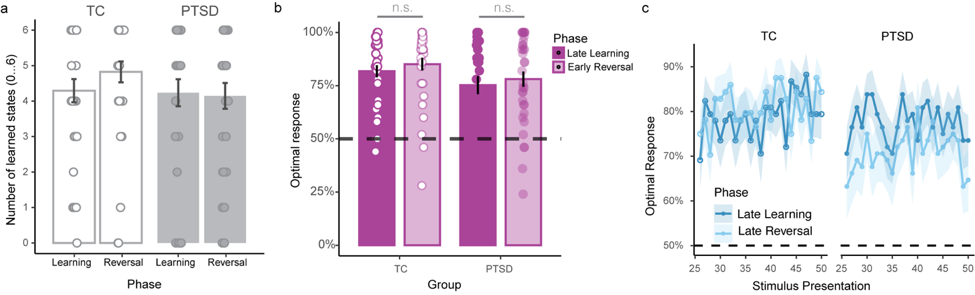

**Supplementary Figure 3. Whole-brain analysis of learning.** **a.** Whole-brain contrasts of Late versus Early Learning, collapsed across groups. Yellow boxes indicate the striatum. Color bars indicate z-static range. **b.** Group comparison of Late versus Early Learning. Yellow box indicates a significant cluster in the putamen. **c.** Whole-brain contrast of optimal versus non-optimal responses, showing significant cluster in striatum (putamen and caudate). All results shown are voxel-wise FDR-corrected at *p* < 0.05.

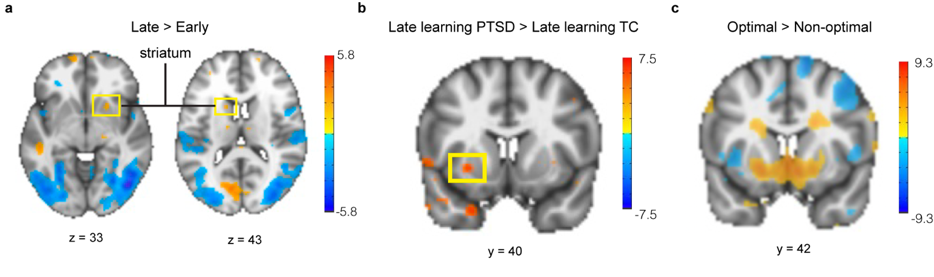

**Supplementary Figure 4. Computational model diagnostics, validation, and parameter-behavior relationships.** **a.** MCMC chain trace plots for each of 4 chains, for each population-level estimated parameter shown across 2000 post-warmup iterations; chains show good mixing and convergence. **b.** Parameter recovery for TC (top) and PTSD (bottom). True parameters from model estimation are correlated with recovered parameter estimates for each model free parameter; all parameters were strongly recovered. **c.** Posterior distributions of groups differences in each of the three parameters. Red bars indicate 95% highest density interval (HDI); all distributions overlap 0. **d.** Parameter-behavior correlations for TC and PTSD groups across Learning, Reversal, and Devaluation. Opt = percent of optimal (80% reinforced) responses; winstay = tendency to repeat rewarded actions for a given stimulus; switch = tendency to switch responses on consecutive trials. Correlation values reflect Pearson’s *r*. Warmer colors indicate positive correlations; cool colors indicate negative correlations. **p* < 0.05

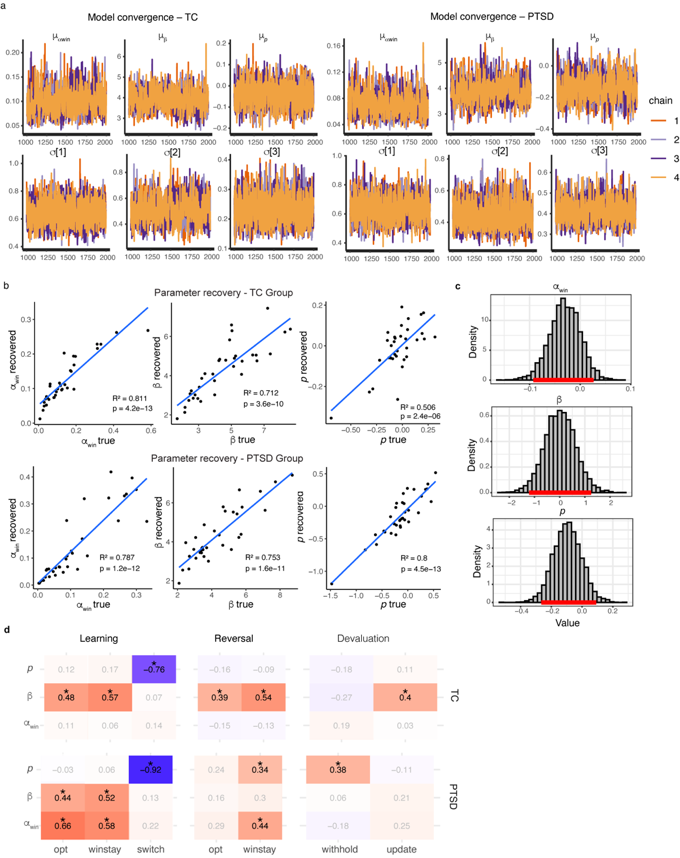

**Supplementary Figure 5. Associations of symptom severity and behavior.** **a.** Correlations of overall symptom severity (PCL score) and adaptive behavior in Learning, and with learning rate. **b.** Exploratory analysis, showing correlations of adaptive behavior in learning with PCL cluster E (Arousal) scores, with learning rate, and with Putamen responses in Late Learning. Dark Lines denote PTSD, light lines denote TC, and shaded regions represent 95% CI.

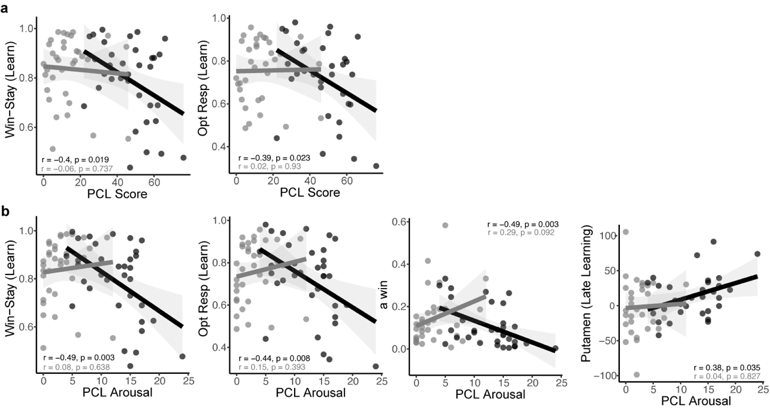
